# Drought adaptation in nature by extensive genetic loss-of-function

**DOI:** 10.1101/372854

**Authors:** J.G. Monroe, T. Powell, N. Price, J.L. Mullen, A. Howard, K. Evans, J.T. Lovell, J.K. McKay

## Abstract

Visions of a second green revolution empowered by emerging technologies have called for interdisciplinary syntheses to scale up the discovery of functionally definitive gene variants responsible for climate adaptation in plants. We integrated novel approaches using whole genome sequences and satellite remote sensing to identify natural knockout alleles associated with drought histories in wild *Arabidopsis thaliana*. Genes identified exhibit signatures of parallel molecular evolution, selection for loss-of-function alleles, and shared associations with flowering time phenotypes in directions consistent with longstanding adaptive hypotheses 7 times more often than expected by chance. Artificial knockout lines then confirmed predicted phenotypes experimentally. These findings further challenge popular assumptions about the adaptive value of genetic loss-of-function in nature and inspire new opportunities for engineering climate resilience in crops.

**One sentence summary:** Whole genome sequences and satellite-detected droughts point to gene knockouts as valuable genetic fuel for climate adaptation.

---

Droughts affect billions of people each year and pose the greatest threat to global food stability. Accelerated efforts to reverse engineer drought tolerance in crops can be informed through insights gained from adaptation in wild plant species but require interdisciplinary perspectives and translatable discoveries (*1*). Such an evolutionary research program is motivated by the need to understand adaptive drought tolerance strategies for different types of drought conditions, which can vary in severity and timing (*2*). Furthermore, previous failures of single locus approaches have reinforced the necessity of developing methods to identify beneficial alleles at both genomic scales and functional molecular resolutions to yield a more complete understanding evolution and discoveries relevant to future molecular breeding decisions (*3, 4*). We combined long-term satellite-detected drought histories, whole genome sequence scans based on allele function, and hypothesis driven reverse genetic screening in *Arabidopsis thaliana* (*Arabidopsis* hereafter) to test historical predictions about how drought timing shapes the evolution of flowering time and outline a broadly scalable approach for discovering functionally definitive gene variants responsible for plant climate adaptation.

Plants have been adapting to drought for millennia and evolutionary responses to specific drought events can be dramatic (*5*). Historically, most research has studied late growing-season droughts, yet drought conditions can occur throughout the year and drought timing is forecast to change over the next century (*6*). Nevertheless, the observation both in nature and agriculture that plants are particularly susceptible to drought while flowering (*7, 8*) has contributed to the longstanding hypothesis that adaptive flowering time evolution should reflect patterns in the seasonal timing of drought events (*9*). Detailed studies of life history (*10*) also reveal that locally adapted *Arabidopsis* populations begin flowering in their home environments just prior to and after periods of increased historical drought frequency (Fig. 1A and B). This motivated an investigation to identify alleles associated with drought timing and address the hypothesis that they contribute to adaptive flowering time evolution.

**Fig. 1.**
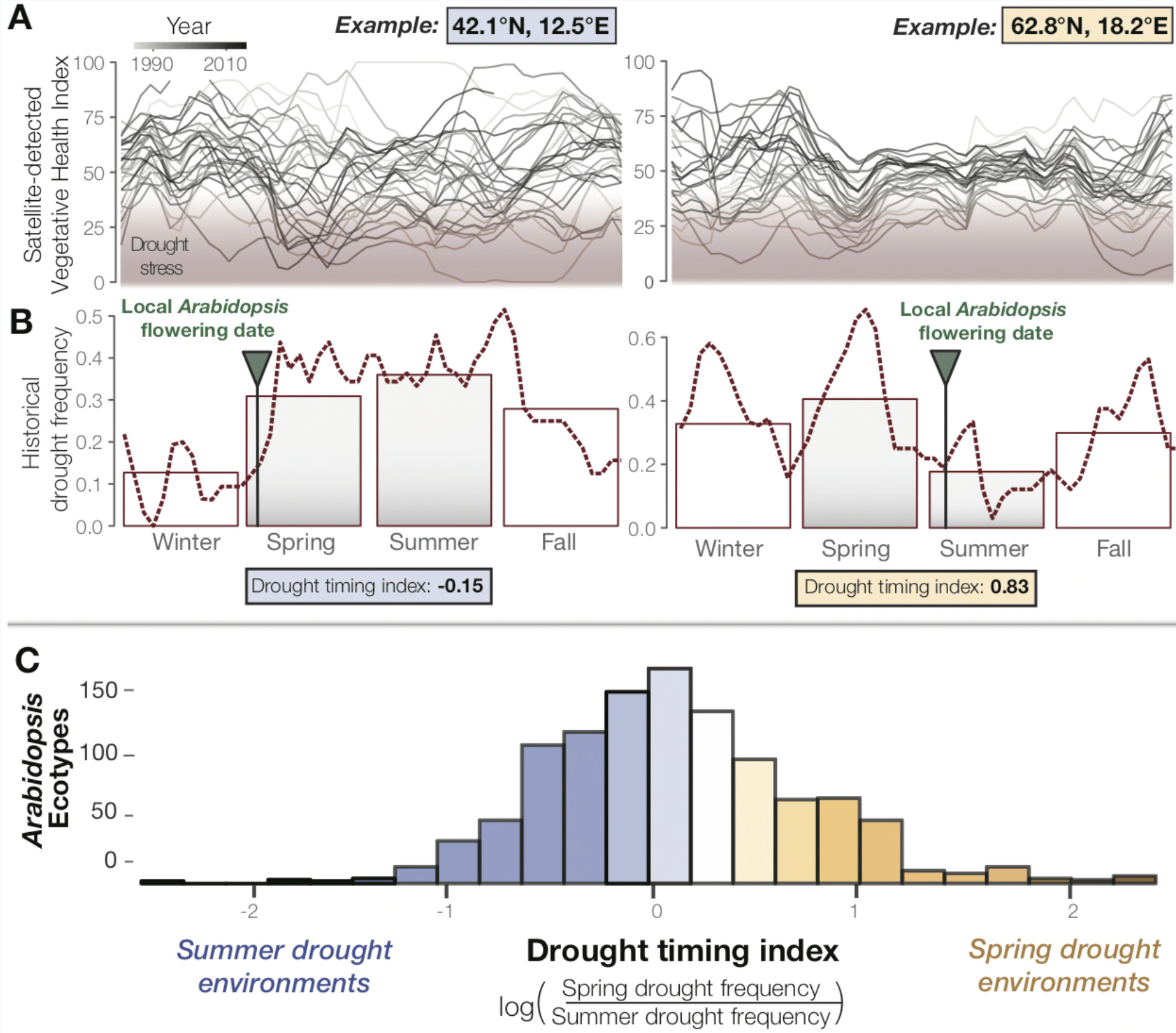
Seasonal drought timing varies across the species range of *Arabidopsis*. (**A**) Examples of home environments for two well-studied *Arabidopsis* ecotypes (*10*) showing historical drought conditions detected using the VHI and (**B**) drought frequency (VHI<40) by week (line) and season (bars). Arrows mark locally observed flowering dates (*10*) and gray bars highlight typical reproductive growing season used to quantify drought-timing index. (**C**) Variation in historical drought timing experienced at the home environments of *Arabidopsis* ecotypes across the much of the species range (fig. S1, table S1). Large values indicate environments where spring droughts occur more frequently than summer drought (i.e. where the frequency of drought decreases over the course of the typical reproductive growing season) and vice versa.

To study global seasonal drought timing, satellite-detected measurements offer a valuable historical record. One such measurement, the Vegetative Health Index (VHI) has been used for decades to predict crop productivity (*11*). By quantifying drought induced vegetative stress this index also presents a resource to study seasonal patterns in drought-related episodes of natural selection. We analyzed 34 years of VHI data to characterize drought regimens at the home environments of *Arabidopsis* ecotypes (e.g. Fig. 1A and B). We then generated a drought-timing index that quantifies the relative frequency of drought at different times over the typical reproductive growing season and observed substantial differences in drought timing experienced by ecotypes (Fig. 1C, Fig. S1, Data S1).

Translating the evolutionary genetics of drought adaptation to crop improvement requires the identification of causative and functionally definitive gene variants. In contrast to early theoretical predictions, loss-of-function (LoF) alleles, those that eliminate or ‘knockout’ a gene’s molecular function, are overrepresented among alleles reported as responsible for crop improvement and often produce adaptive phenotypes in wild species (*12-15*). Adaptive LoF alleles are also particularly valuable for targeted molecular breeding because functionally similar mutations can be mined from the breeding pool or generated directly by non-transgenic native gene editing. Unfortunately, traditional genome-wide association scans relying on the one-locus, two-allele model perform poorly at detecting adaptive LoF alleles, which often arise through parallel molecular evolution (*16, 17*). Species-wide whole genome sequences, however, present the opportunity to advance beyond previous mapping and scanning methods by contrasting predicted functional allele states rather than SNPs, to understand evolutionary genetics at functional molecular resolutions and inform perspectives on next generation molecular breeding.

We analyzed whole genome sequences in *Arabidopsis* to identify candidate LoF alleles underlying drought adaptation and flowering time evolution. We first surveyed the genomes of 1135 ecotypes (*18*) for LoF alleles in protein coding genes predicted to encode truncated amino acid sequences (Data S2). To overcome the potential parallel evolutionary origins of LoF alleles that would have challenged previous methods, we classified alleles based functional allele state rather than individual polymorphisms for association testing. After filtering steps to reduce the likelihood of false positives, we thus tested 2088 genes for LoF allele associations with drought timing (Fig. 2A) and flowering time (Fig. 2B). These analyses identified 247 genes in which LoF alleles are significantly associated with drought timing and/or flowering time after accounting for population structure and multiple testing (Data S3).

**Fig. 2.**
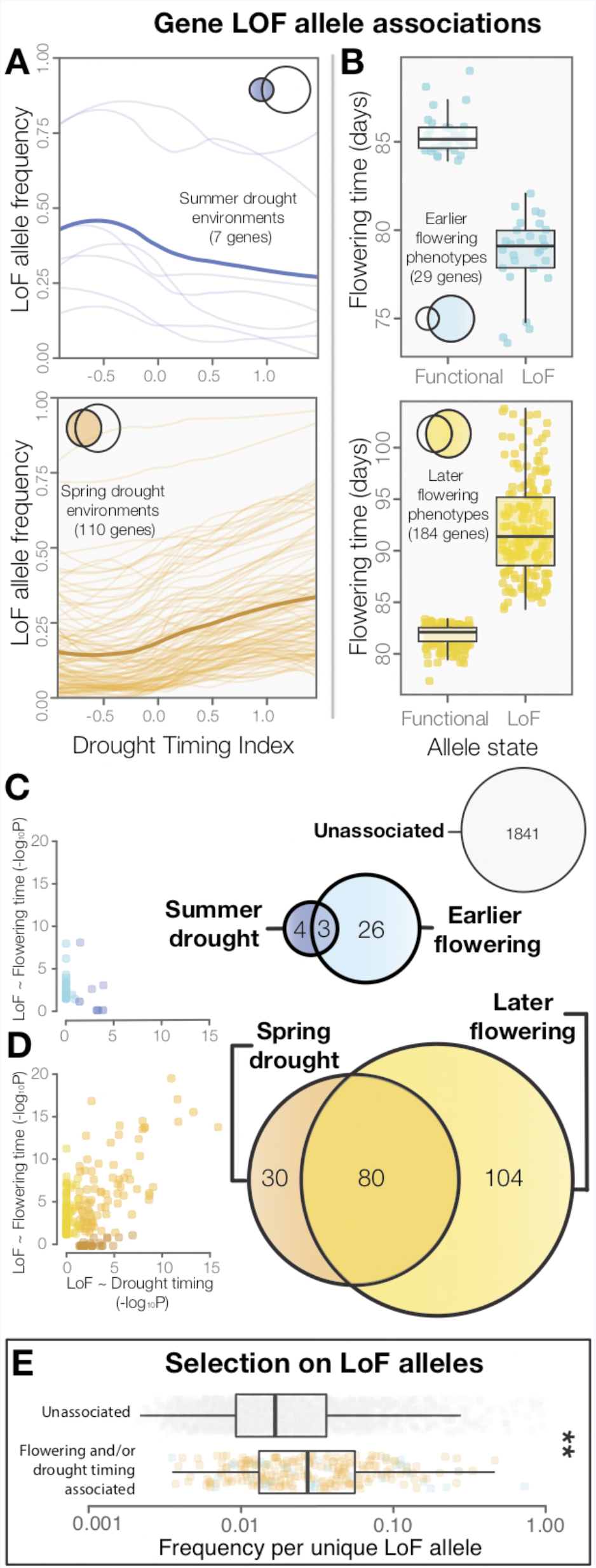
LoF alleles share associations between drought timing and flowering time, and show evidence of positive selection. (**A**) Visualization of the frequency of LoF alleles across environments in genes associated to summer (upper) or spring drought environments (lower). Darker lines indicate the mean across genes. (**B**) Contrasting flowering times between ecotypes with functional versus LoF alleles in genes associated with earlier (upper) or later (lower) flowering time phenotypes. (**C**) Overlap and relationships between the strength of LoF allele associations in genes associated with summer drought and earlier flowering, and (**D**) spring drought and later flowering. (**E**) Increased frequencies of independent LoF alleles in genes associated with drought timing and/or flowering time compared to genes without detected associations (t-test, P=3.4×10^-7^), a signature of recurrent mutation accompanied by positive selection (*16*).

Associations to drought timing predicted associations of LoF alleles to flowering time directly. Together, summer drought and earlier flowering associated genes (Fig. 2C), and spring drought and later flowering associated genes (Fig. 2D) overlapped 7 times more often than expected by chance (χ^2^=492, P<2 ×10^-16^) and no shared associations were observed in the opposite direction. This result supports the classical hypothesis that flowering time reflects adaptation to local drought regimens, indicating the evolution of “escape” through earlier flowering in summer drought environments, and “avoidance” by later flowering in spring drought environments (*19*). Satellite-detected drought histories thus prove useful for predicting the direction and molecular targets of phenotypic evolution, identifying immediate candidates for hypothesis testing and potential use in developing climate adapted crop varieties. Functional genome-wide association scans with ecologically meaningful environmental variation could be valuable for discovering candidates underlying other important traits that are especially difficult to measure.

Signatures of selection in the genes identified differ from the genome average and neutral expectations. As expected for genes harboring LoF alleles, these genes show parallel evolution of LoF and accelerated amino acid sequence evolution among *Arabidopsis* ecotypes (Fig. S2A and B, Data S4). We also found evidence of positive selection for LoF alleles in genes associated with drought timing and/or flowering time. While these genes have similar global frequencies of LoF alleles compared to genes not showing associations with drought timing and/or flowering time (Fig. SC), they tend to have significantly fewer unique LoF alleles (Fig. 2D) and greater frequencies of each independent LoF allele (Fig. 2E). This pattern is consistent with theoretical predictions and results from simulations of adaptation by parallel molecular evolution involving recurrent mutation combined with more rapid local fixation of alleles experiencing positive selection (*16*).

The extent of LoF responsible for adaptive phenotypic evolution is much greater than once assumed (*20, 21*). LoF alleles identified were overwhelmingly associated with spring drought/later flowering rather than summer drought/earlier flowering (χ^2^ = 132, P < 2×10^-16^, Fig. 2). Because the reference genome and gene models are from an early flowering *Arabidopsis* line, this is consistent with the hypothesis that LoF alleles are particularly important in the evolution of phenotypic divergence (*13*). We found that flowering time is strongly predicted by the accumulation of LoF alleles across the 214 candidate genes associated to spring drought and/or later flowering time (Fig. 3A-E), estimating a 1-day increase for every 3 additional LoF alleles across these candidate genes (Fig. 3F). Importantly, we did not find a broader overabundance of LoF alleles in later flowering ecotypes or those from spring drought environments that would explain this relationship (Fig. S3). Rather, these findings support a model of climate-associated evolution in complex traits that includes a substantial contribution from widespread genetic LoF and give promise to targeted LoF as useful for directed phenotypic engineering.

**Fig. 3.**
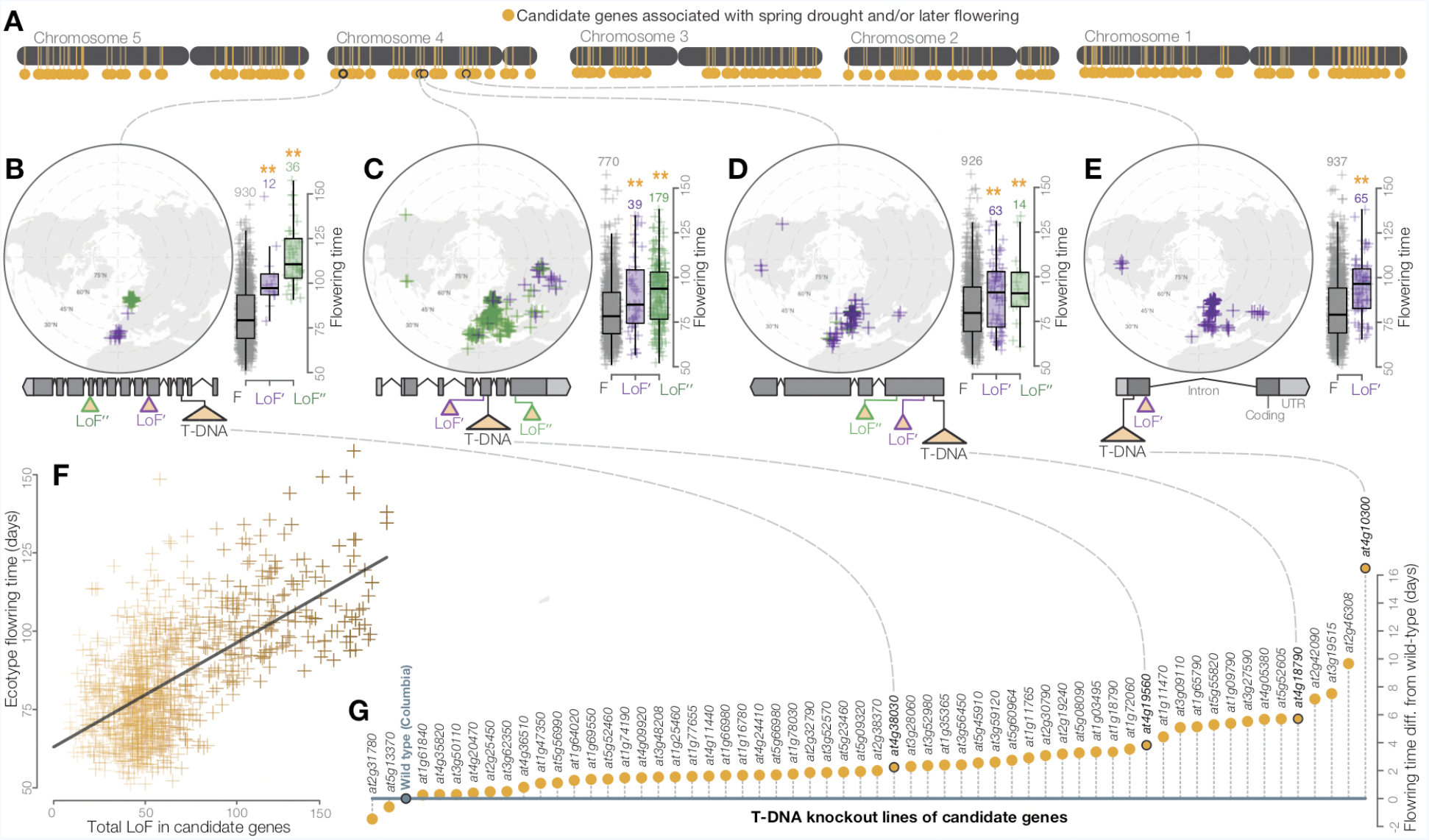
Widespread LoF contributing to later flowering time evolution. (**A**) Genomic map of 214 candidate genes with associations between LoF alleles and spring drought environments and/or later flowering time phenotypes. (**B-E**) Examples of the geography and flowering times among *Arabidopsis* ecotypes of LoF alleles in candidate genes including; (**B**) a previously unstudied rhamnogalacturonate lyase, (**C**) a cyclin linked to later flowering in prior knockout experiments (*22*), (**D**) members of the drought-responsive Nramp2 (*23*) (**E**) and RmlC-like cupin (*24*) protein families. (**F**) Later flowering time in ecotypes predicted by the accumulation of LoF alleles across all candidate genes. Color scale of points reflects proportion of total LoF in ecotypes that are candidate genes (darker points = greater proportion) (**G**) Experimental validation of hypothesized later flowering time in T-DNA knockout lines of candidate genes compared to the wild type genotype.

Experimental knockout lines confirmed the later flowering times predicted from natural allele associations. To test phenotypic effects we screened a panel of confirmed T-DNA insertion mutants representing a sample of candidate LoF alleles associated with spring drought and/or later flowering. As predicted by variation among *Arabidopsis* ecotypes (Fig. 2D), the vast majority of knockout lines in these candidate genes (57 of 59) flowered later on average than the wild type genotype (Fig. 3G, Data S5). LoF alleles identified through these analyses and experiments include those previously linked to flowering time (*22*) and drought responses (*23, 24*). Implementing a functional genome-wide association scan, we find that allele associations with ecologically meaningful environmental variation (drought timing) accurately predict associations with adaptive phenotypes directly (flowering time). Together with validation in transgenic lines, these findings outline a scalable model for gaining deeper insights into the functional genomics of climate adaptation in nature and further challenge historical assumptions about molecular adaptation that have implications for influencing evolutionary theory and public attitudes toward emerging molecular breeding approaches.

Groundbreaking yield increases during the green revolution of the 1960s were largely attributable to semi-dwarf phenotypes caused by LoF alleles in both rice and barley (*25, 26*). Later it was found that natural LoF alleles of the same gene in wild *Arabidopsis* produce similar phenotypes (*17*), suggesting the potential to mine ecological species for information directly useful for crop improvement. Visions of a second green revolution powered and informed by such natural variation call for discoveries in evolutionary functional genomics at scales that have now become possible. The work presented here demonstrates a critical step in this process of exploiting evolution to inspire future molecular breeding of climate resilient crops and highlights the value of integrating breakthrough technologies from diverse disciplines to develop a more complete understanding of evolution.

## Acknowledgments

E. Buckler, D. Des Marais, A. Henry, J. Lasky, T. Mitchell-Olds, J. Ross-Ibarra, and D. Sloan provided valuable feedback and insightful discussion that improved this work. This study was financially supported by NSF Awards DEB 1022196 and 1556262 to JKM, NSF Award 1701918 and USDA-NIFA Award 2014-38420-21801 to JGM, as well as generous funding from Cargill, Inc. Data used are included in the main text, supplementary materials, and public repositories.

## Supplementary Materials

### Materials and Methods

#### Satellite-Detected Drought Histories of *Arabidopsis*

To study patterns in historical drought, the remotely sensed Vegetative Health Index (VHI) was used, a satellite-detected drought measurement tool whose advantage is that it includes information about vegetative impacts of drought (*9, 27*). This index is based on multiple data sources from NOAA satellites, combining deviations from historic climatic (Temperature Condition Index derived from AVHRR-based observations in thermal bands) and vegetative conditions (Vegetative Condition Index derived from NDVI) to detect periods of ecological drought conditions and distinguish between other sources of vegetative stress such as cold (*11, 28, 29*). VHI was collected weekly since 1981 at 16 km^2^ resolution on a scale from 0 to 100, where values below 40 reflect drought conditions (*11*) (Fig. 1A). The frequencies of observing drought conditions during photoperiodic spring (quarter surrounding spring equinox), summer (quarter surrounding summer solstice), fall (quarter surrounding fall equinox), and winter (quarter surrounding winter solstice) were calculated globally from 1981 to 2015 (Fig. 1B) in R (*30*) using the *raster* package (*31*).

To characterize the seasonal timing of droughts during an important period of *Arabidopsis’* life history, a univariate drought-timing index was generated that quantifies whether the historical frequency of drought increases or decreases over the course of the typical *Arabidopsis* reproductive growing season (*32-34*). Specifically, this index is equal to the natural log transformed ratio between spring and summer drought frequency. More negative values reflect environments where drought frequency increases from spring to summer and are referred to here as “summer drought environments,” (e.g. Fig. 1B left). Conversely, more positive values reflect environments where drought frequency decreases from spring to summer and are referred to here as ‘spring drought environments,’ (e.g. Fig. 1B right). After removing ecotypes with missing location data or locations falling within pixels classified as water, seasonal drought frequencies and drought timing were calculated at the location of origin for 1,097 *Arabidopsis* ecotypes that were included as part of the 1001 Genomes Project (*18*)(Fig. 1C, table S1). Up to date global map files of seasonal drought frequency and the drought-timing index used here are available on Dryad and greymonroe.github.io/data/drought alongside a brief tutorial showing how to extract data for points of interest in R.

#### Loss-of-Function (LoF) Alleles in *Arabidopsis* Genomes

To identify functionally definitive gene variants (*35-37*), LoF alleles (*21*) were identified from whole genome sequence data of 1,135 *Arabidopsis* accessions (*18, 38,39*) using R scripts available at greymonroe.github.io/data. First, genes were filtered to those containing at least 5% frequency of predicted frameshift or premature stop mutations from results generated by the 1,001 Genomes Consortium (*18*) using ‘*SnpEff’* (*40*). To reduce instances where exon skipping might ameliorate LoF mutations (*41*), genes were filtered to those with a single predicted gene model (*42*). Additionally, to preclude false LoF calls for cases where compensatory mutations restore gene function or in which an insignificant portion of the final protein product is affected by putative LoF mutations (*43*), coding regions were translated into predicted amino acid sequences from which lengths from start to stop codon were calculated (table S2) using *seqinr*. LoF alleles were defined as those producing protein products with at least 10% lost because of late start codons and/or prematurely truncated translation. Allelic heterogeneity expected to mask these genes from traditional GWAS (*44-46*) was corrected for by classifying all alleles as either functional (0) or non-functional (1) (table S3). A final frequency filter was re-applied (5% global LoF allele frequency), resulting in 2088 genes for downstream association analyses.

#### LoF Associations to Drought Timing and Flowering Time

To identify candidate LoF alleles responsible for climate adaptation and phenotypic evolution, the relationships between functional allele state and drought timing and between functional allele state and flowering time were evaluated for each of the 2088 genes that passed preceding filtering steps. Specifically, the association between functional allele state among *Arabidopsis* ecotypes and historical drought timing at their locations of origin was tested by logistic regression in a generalized linear model in R (*30*). To reduce false positives and false negatives, population structure was accounted for by performing a principal component analysis on the kinship matrix among all ecotypes (*18*) and including in each model the first three resulting principal components, which explain >75% of variance in relatedness between ecotypes (*47*). The P-values (P_drought timing_) of the slope estimates (β_drought timing_) for drought timing in these models were adjusted to account for multiple tests by a Bonferroni correction (table S4).

Summer drought genes were identified as those in which LoF alleles are found in ecotypes that experience a significantly (β_drought timing_ <0 & _drought timing_ <0.05) more negative drought-timing index (summer drought environments where drought frequency increases over the course of the reproductive growing season, Fig. 1B left and Fig. 2A top). Conversely, spring drought genes were identified as those in which LoF alleles are found in ecotypes that experience a significantly (β_drought timing_ > 0 & P_drought timing_ <0.05) more positive drought-timing index (spring drought environments where drought frequency decreases over the course of the reproductive growing season, Fig. 1B right and Fig. 2A bottom).

The above analytical approach was repeated to test whether functional allele state is associated with the reported common garden flowering times of *Arabidopsis* ecotypes (*14*) (table S4). See Alonso-Blanco *et al.* (*14*) for details, but in brief, flowering time was measured in growth chambers at 10°C (considerably less missing data than experiment at 16°C) under 16 hour days. Earlier flowering genes were identified as those in which LoF alleles are found in ecotypes that flower significantly (β_flowering time_ <0 & P_flowering time_ <0.05) earlier than ecotypes with a functional allele (Fig. 2B top). Later flowering genes were identified as those in which LoF alleles are found in ecotypes that flower significantly (β_flowering time_ >0 & P_flowering time_ <0.05) later than ecotypes with a functional allele (Fig. 2B bottom).

#### Overlap Between Drought Timing and Flowering Time Associated Genes

To address the longstanding hypothesis that flowering time reflects adaptation to drought timing (*9, 19, 48*), and the test the corresponding prediction that alleles associated with drought timing are also associated with flowering time, the groups of genes identified with significant associations to drought timing or flowering time were compared (Fig. 2C and D). Deviation from the null hypothesis of independent associations to drought timing and flowering time was evaluated by a chi-squared test (Expected number of co-associated genes = 12, Observed = 83, χ^2^=492, P=2×10^-16^).

The magnitude of P-values have historically served as the basis of selecting candidate loci for further examination toward their contribution to environmental adaptation or phenotypic evolution in quantitative trait locus mapping and genome wide association scans [e.g. (*49*)]. To test whether associations to environment (drought timing) can be used to identify loci associated with phenotypes (flowering time) directly, the correlation between log transformed P-values describing allele associations with drought timing (P_drought timing_) and with flowering time (P_flowering time_) was calculated (r^2^=0.52, P<2×10^-16^) and visualized separately for genes associated to summer drought/earlier flowering (Fig. 2C) and to spring drought/later flowering (Fig. 2D).

#### Signatures of Selection

To assess whether histories of selection for genes identified differ from the genome wide expectation, measures of amino acid sequence evolution were evaluated for 122 genes in which loss-of-function is associated with drought timing or flowering time and for which there are orthologs identified between *A. lyrata* and *A. thaliana* (*50*). For each gene, sequences were aligned using MAFFT (*51*), codons with gaps removed, and the number of non-synonymous and synonymous polymorphisms among *A. thaliana* accessions (P_N_ and P_S_) as well as synonymous and non-synonymous divergence (D_N_ and D_S_) from *A. lyrata* were measured using mkTest.rb (https://github.com/kern-lab/). The ratios P_N_/P_S_ and D_N_/D_S_ were then calculated to measure the proportion of variants predicted to affect amino acid sequences that are segregating among ecotypes and diverged from *A. lyrata*, respectively. These calculations were also performed for genes not associated to drought timing or flowering time (n=912) and the remaining genes across the *A. thaliana* genome (n=20373) with orthologs between *A. lyrata* and *A. thaliana*. To test whether genes identified show evidence of accelerated protein sequence evolution, comparisons were made to genes associated with drought timing or flowering time for both P_N_/P_S_ (Fig. S1A) and D_N_/D_S_ (Fig. S1B) by two-sided students t-tests (α=0.05) in R (*30*).

Because theory predicts adaptation by loss-of-function to proceed through multiple independent alleles, but to exhibit a fewer number of different alleles than in neutral loci at similar LoF allele frequencies (*16, 52, 53*), the number of unique LoF alleles was estimated by protein length in the genes that passed preceding filtering steps. To address the hypothesis that genes in which LoF alleles are associated to drought history or flowering time are likely to reflect positive selection compared to genes in which LoF are random with respect to drought history or flowering time, the total number of unique LoF alleles between these groups was compared using a two-sided students t-test (log10 transformed, P=5.8×10^-7^, Fig. S2D). To control for the possibility that this result in an artifact of reduced frequency of LoF alleles in genes identified, the global frequency of LoF was also compared between these groups (log_10_ transformed, two-sided students t-test, P=0.11, Fig. S2C). Finally, to further test the prediction that LoF alleles in genes identified have increased in frequency because of more positive selection, the frequency per specific LoF allele was compared between groups (log_10_ transformed, two-sided students t-test, P= 3.4×10^-7^, Fig. 2E).

#### Candidate Genes Contributing to Later Flowering Time by Widespread LoF

The significance of the tendency for LoF associations to spring drought/later flowering time (Fig. 2D) was tested by X-squared tests (spring drought vs. summer drought, P<2×10^-16^; later vs. earlier flowering, P<2×10^-16^, spring drought/later flowering vs. summer drought/earlier flowering, P<2×10^-16^). The chromosomal locations of candidate genes (those associated to spring drought/later flowering time) were mapped onto the *Arabidopsis* genome (*42*) (Fig. 3A). To address the hypothesis that widespread LoF contributes to later flowering time phenotypes, the total number of LoF in candidate genes for each ecotype was calculated and the correlation between this value and flowering time evaluated (Fig. 3F, r^2^=0.39, P<2×10^-16^).

#### Experimental Testing of Predicted Phenotypes in Gene Knockout Lines

The preceding analyses provided compelling evidence of LoF in candidate genes as important in the evolution of later flowering time phenotypes. To test the prediction that non-functionalization of these genes causes increased flowering time, phenotypes were measured in transgenic lines in a subsample of candidate genes showing a significant association between loss-of-function and spring drought environments and/or later flowering time. Motivated by the general need to develop a high throughput approach of studying naturally adaptive LoF, knockout lines from the Arabidopsis Biological Resource Center were chosen from a collection created by the SALK Institute in which a T-DNA insertion in an exon of candidate genes has already been identified and confirmed to be homozygous (*54, 55*). These T-DNA knockout lines were generated by the SALK institute (Data S5) and exist in a common genetic background (Columbia) (*56*). Seeds were planted in 2” pots containing wet potting soil and stratified for 5 days at 4°C. Seedlings were thinned to a single plant per pot one week after stratification. Plants were grown (59 T-DNA knockout lines, 10 reps of each line and 30 reps Columbia) in a stratified (by shelf), randomized design in growth chambers (Conviron ATC60, Controlled Environments, Winnipeg, MB) under 16 hours of light at 20°C. Flowering time was measured as days after planting to the emergence of the first open flower. We calculated the least squares mean (lsmean from ‘lsmeans’ package in R) flowering time for each line from a mixed model where shelf and tray were included as random effects (table S5). We tested the prediction that knockout lines would flower later (have higher lsmean flowering time estimates) than the wild type Columbia genotype by a X-squared test (P=8.1×10^-13^).

---

**Fig. S1.**
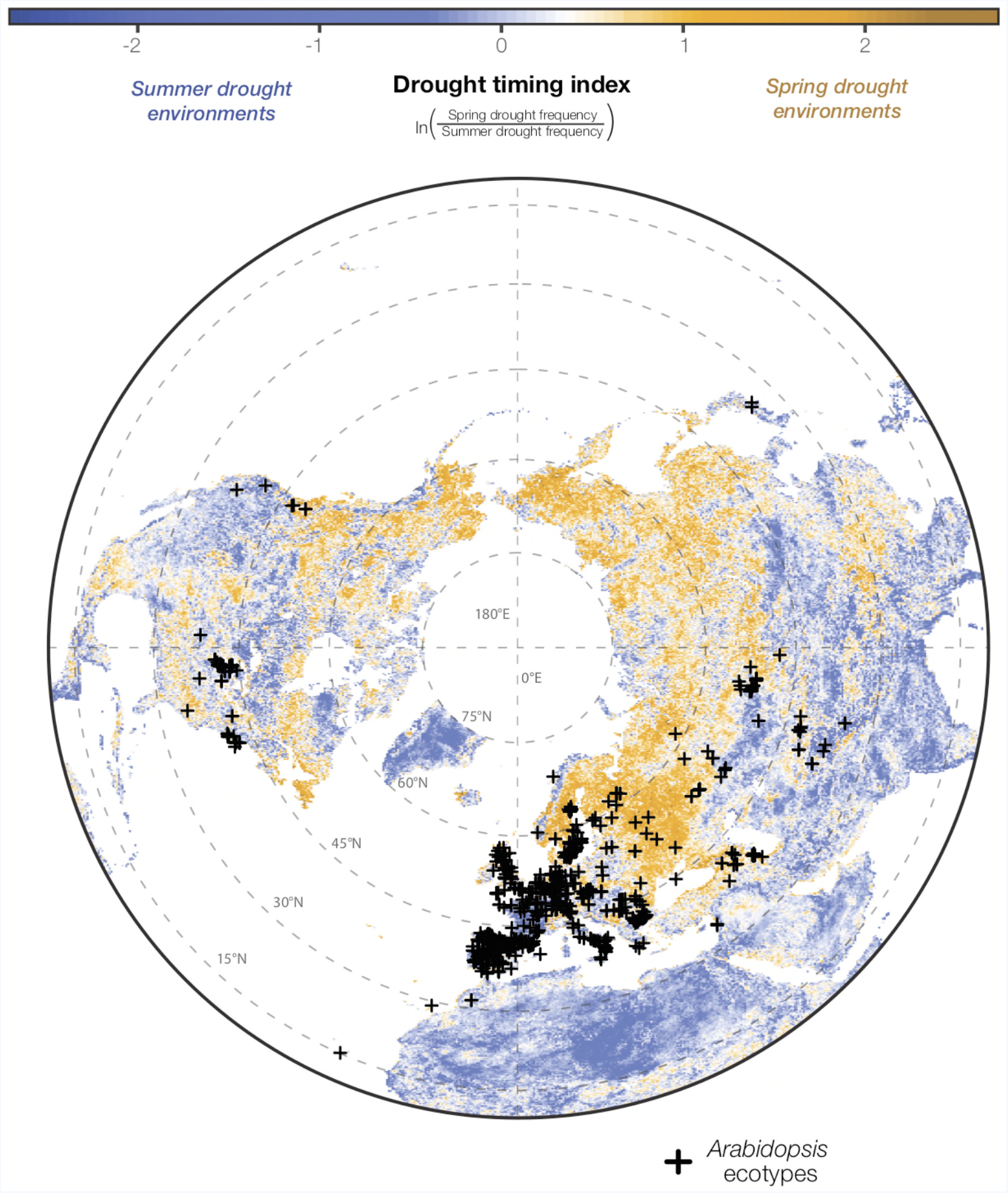
*Arabidopsis* ecotypes are distributed across satellite-detected drought timing gradients. Historical patterns in drought conditions were calculated from the Vegetative Health Index (VHI, Fig. 1A) and converted into a drought-timing index (Fig. 1B and C). Large values of this index indicate environments where spring droughts occur more frequently than summer drought (ie. where the frequency of drought decreases over the course of the reproductive growing season) and vice versa (raw map data available at greymonroe.github.io/data/drought).

**Fig. S2.**
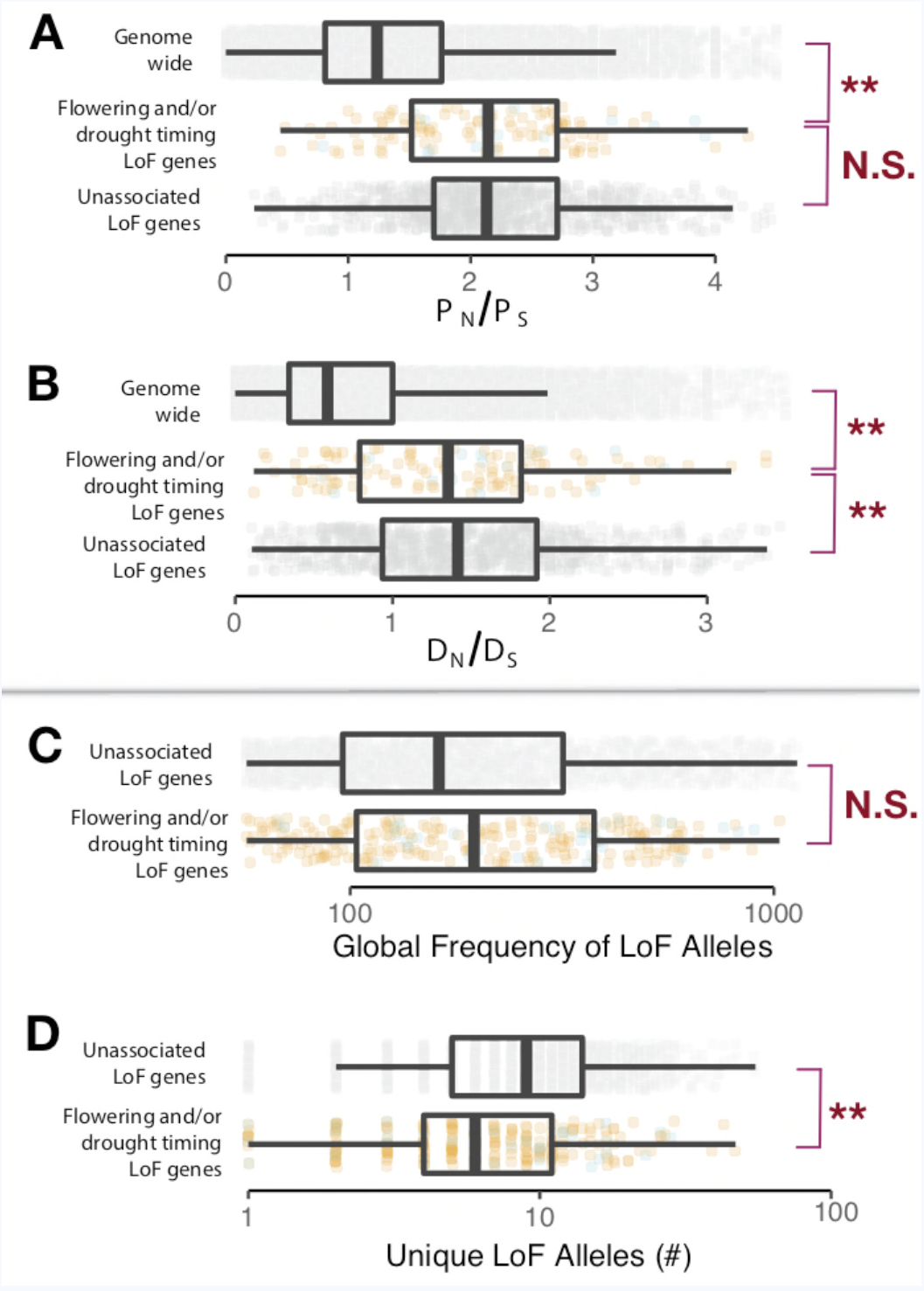
Signatures of selection on LoF genes identified differ from null expectations. (**A**) Contrasts (t-test, α=0.05) between genes identified with LoF alleles associated to drought timing and/or flowering time (colors correspond to Fig. 2 C and D, boxplots visualized at ±1.5 times the data interquartile range) and the genomic background (light gray), as well as genes having LoF alleles but without observed associations (dark gray) for the ratio of non-synonymous (P_N_) and synonymous polymorphisms (P_S_) among *A. thaliana* ecotypes and (**B**) the ratio of non-synonymous (D_N_) and synonymous divergence (D_S_) from *A. lyrata*. (**C**) Contrasts (t-test, α=0.05) between (log_10_) global frequency of LoF alleles in genes identified with LoF alleles associated to drought timing and/or flowering time and genes with LoF alleles but without observed associations for the global frequency of LoF alleles and (**D**) the number of (log_10_) unique LoF alleles. The corresponding average frequencies of unique LoF alleles for genes are shown in Fig. 2E.

**Fig. S3.**
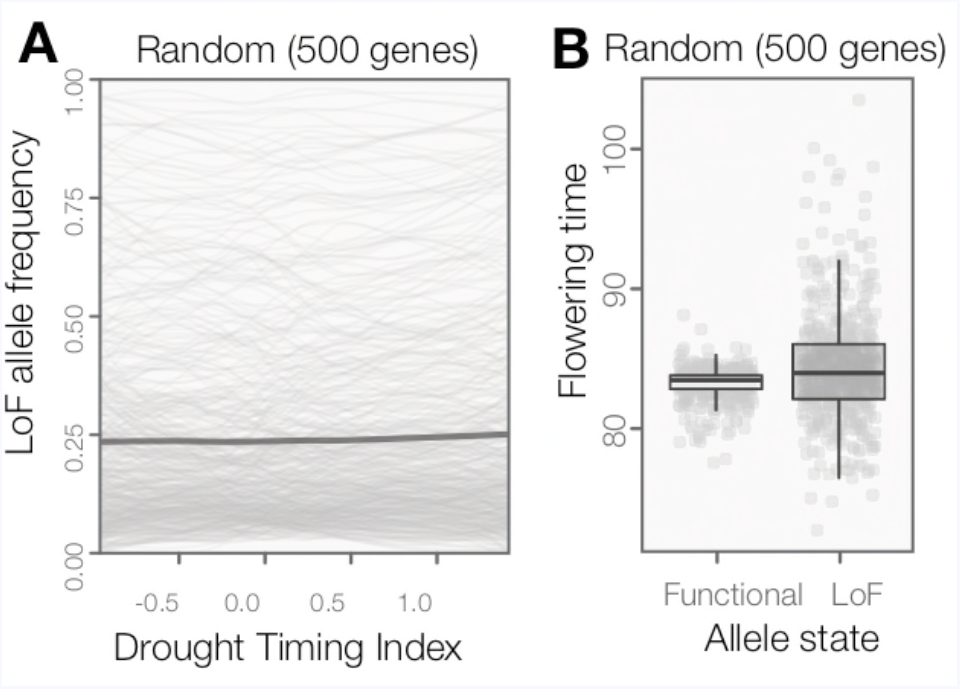
LoF alleles are not broadly overabundant in *Arabidopsis* ecotypes originating from spring drought environments or flowering later. (**A**) The frequency of LoF alleles across environments (sliding window plot) in random genes. The darker line indicates the mean across genes. The distribution of LoF alleles in these random genes contrasts with LoF alleles in genes associated to drought timing, which are overwhelming associated to spring drought environments (Fig. 2A) (**B**) Flowering times compared between ecotypes with functional versus LoF alleles in random genes. The phenotypic differences predicted by these random genes contrasts with LoF alleles in those associated to flowering time, which are overwhelming associated to later flowering time (Fig. 2B)

**Data S1. *Arabidopsis* ecotypes examined.**

Includes ecotype identifiers as well as latitude and longitude of origin, seasonal drought frequencies (winter, spring, summer, fall), drought timing index (drought_timing), and flowering time (FT10).

**Data S2. Matrix of functional allele calls for 2088 genes among 1135 *Arabidopsis* ecotypes.**

LoF alleles are those with less than 90% predicted protein product and are classified with a “1”. Function alleles are classified with a “0”.

**Data S3. Associations between functional allele state and drought timing and flowering time for 2088 genes.**

Includes gene, estimate for logistic regression model testing the association between functional allele state and drought timing (DT_B) and flowering time (FLW_B) after accounting for population structure, and the P-value of these estimates after Bonferroni correction for multiple testing (DT_p and FLW_p)

**Data S4. Selection statistics for 2088 genes.**

Includes P_N_/P_S_ (pnps), D_N_/D_S_ (dnds), frequency, number of LoF alleles, and average frequency per LoF allele.

**Data S5. Flowering time in T-DNA knockout lines.**

Flowering time (lsmean and standard error) of wild-type genomic background and T-DNA knockout lines of a sample of candidate genes in which LoF alleles are associated with spring drought environments or later flowering time phenotypes in *Arabidopsis* ecotypes.

## References and Notes

1. M. V. Mickelbart, P. M. Hasegawa, J. Bailey-Serres, Genetic mechanisms of abiotic stress tolerance that translate to crop yield stability. Nat Rev Genet 16, 237–251 (2015).

2. F. Tardieu, Any trait or trait-related allele can confer drought tolerance: just design the right drought scenario. Journal of experimental botany 63, 25–31 (2011).

3. J. Passioura, Scaling up: the essence of effective agricultural research. Functional Plant Biology 37, 585–591 (2010).

4. A. M. Dean, J. W. Thornton, Mechanistic approaches to the study of evolution: the functional synthesis. Nature Reviews Genetics 8, 675 (2007).

5. S. J. Franks, S. Sim, A. E. Weis, Rapid evolution of flowering time by an annual plant in response to a climate fluctuation. Proceedings of the National Academy of Sciences 104, 1278–1282 (2007).

6. K. E. Trenberth et al., Global warming and changes in drought. Nature Climate Change 4, 17 (2014).

7. N. Nam, Y. Chauhan, C. Johansen, Effect of timing of drought stress on growth and grain yield of extra-short-duration pigeonpea lines. The Journal of Agricultural Science 136, 179–189 (2001).

8. J. D. Dietrich, M. D. Smith, The effect of timing of growing season drought on flowering of a dominant C4 grass. Oecologia 181, 391–399 (2016).

9. J. Passioura, Drought and drought tolerance. Plant Growth Regulation 20, 79–83 (1996).

10. J. P. Mojica et al., Genetics of water use physiology in locally adapted Arabidopsis thaliana. Plant Sci 251, 12–22 (2016).

11. F. N. Kogan, Global drought watch from space. Bulletin of the American Meteorological Society 78, 621–636 (1997).

12. H. E. Hoekstra, R. J. Hirschmann, R. A. Bundey, P. A. Insel, J. P. Crossland, A single amino acid mutation contributes to adaptive beach mouse color pattern. Science 313, 101–104 (2006).

13. M. D. Rausher, Evolutionary transitions in floral color. International Journal of Plant Sciences 169, 7–21 (2008).

14. C. Alonso-Blanco, B. Méndez-Vigo, Genetic architecture of naturally occurring quantitative traits in plants: an updated synthesis. Current opinion in plant biology 18, 37–43 (2014).

15. K. M. Olsen, J. F. Wendel, A bountiful harvest: genomic insights into crop domestication phenotypes. Annu Rev Plant Biol 64, 47–70 (2013).

16. P. Pennings, J. Hermisson, Soft Sweeps III - The signature of positive selection from recurrent mutation. PLoS Genetics preprint, (2005).

17. L. Barboza et al., Arabidopsis semidwarfs evolved from independent mutations in GA20ox1, ortholog to green revolution dwarf alleles in rice and barley. Proc Natl Acad Sci U S A 110, 15818–15823 (2013).

18. C. Alonso-Blanco et al., 1,135 genomes reveal the global pattern of polymorphism in Arabidopsis thaliana. Cell 166, 481–491 (2016).

19. N. J. Kooyers, The evolution of drought escape and avoidance in natural herbaceous populations. Plant Sci 234, 155–162 (2015).

20. J. M. Smith, Natural selection and the concept of a protein space. Nature 225, 563 (1970).

21. R. Albalat, C. Canestro, Evolution by gene loss. Nat Rev Genet 17, 379–391 (2016).

22. X. Cui, B. Fan, J. Scholz, Z. Chen, Roles of Arabidopsis cyclin-dependent kinase C complexes in cauliflower mosaic virus infection, plant growth, and development. The Plant Cell 19, 1388–1402 (2007).

23. L. Qin et al., Genome-Wide Identification and Expression Analysis of NRAMP Family Genes in Soybean (Glycine Max L.). Frontiers in plant science 8, 1436 (2017).

24. M. Aghdasi, F. Fazli, M. B. Bagherieh, Cloning and expression analysis of Arabidopsis TRR14 gene under salt and drought stress. Journal of Cell and Molecular Research 4, 1–10 (2012).

25. W. Spielmeyer, M. H. Ellis, P. M. Chandler, Semidwarf (sd-1),”green revolution” rice, contains a defective gibberellin 20-oxidase gene. Proceedings of the National Academy of Sciences 99, 9043–9048 (2002).

26. Q. Jia et al., GA-20 oxidase as a candidate for the semidwarf gene sdw1/denso in barley. Functional & integrative genomics 9, 255–262 (2009).

27. A. AghaKouchak et al., Remote sensing of drought: Progress, challenges and opportunities. Reviews of Geophysics 53, 452–480 (2015).

28. F. Kogan, B. Yang, G. Wei, P. Zhiyuan, J. Xianfeng, Modelling corn production in China using AVHRR-based vegetation health indices. International Journal of Remote Sensing 26, 2325–2336 (2005).

29. O. Rojas, A. Vrieling, F. Rembold, Assessing drought probability for agricultural areas in Africa with coarse resolution remote sensing imagery. Remote Sensing of Environment 115, 343–352 (2011).

30. T. R. C. Team. (Vienna, Austria, 2017).

31. R. J. Hijmans. (2016).

32. L. T. Burghardt, C. J. Metcalf, A. M. Wilczek, J. Schmitt, K. Donohue, Modeling the influence of genetic and environmental variation on the expression of plant life cycles across landscapes. Am Nat 185, 212–227 (2015).

33. D. Ratcliffe, Adaptation to habitat in a group of annual plants. The Journal of Ecology, 187–203 (1961).

34. L. Thompson, The spatiotemporal effects of nitrogen and litter on the population dynamics of Arabidopsis thaliana. Journal of Ecology, 63–68 (1994).

35. H. E. Hoekstra, J. A. Coyne, The locus of evolution: evo devo and the genetics of adaptation. Evolution 61, 995–1016 (2007).

36. D. Weigel, M. Nordborg, Population genomics for understanding adaptation in wild plant species. Annual review of genetics 49, 315–338 (2015).

37. K. J. Byers, S. Xu, P. M. Schluter, Molecular mechanisms of adaptation and speciation: why do we need an integrative approach? Mol Ecol 26, 277–290 (2017).

38. M. V. Olson, When less is more: gene loss as an engine of evolutionary change. Am J Hum Genet 64, 18–23 (1999).

39. A. D. Cutter, R. Jovelin, When natural selection gives gene function the cold shoulder. Bioessays 37, 1169–1173 (2015).

40. P. Cingolani et al., A program for annotating and predicting the effects of single nucleotide polymorphisms, SnpEff: SNPs in the genome of Drosophila melanogaster strain w1118; iso-2; iso-3. Fly 6, 80–92 (2012).

41. X. Gan et al., Multiple reference genomes and transcriptomes for Arabidopsis thaliana. Nature 477, 419–423 (2011).

42. P. Lamesch et al., The Arabidopsis Information Resource (TAIR): improved gene annotation and new tools. Nucleic acids research 40, D1202–D1210 (2011).

43. D. G. MacArthur et al., A systematic survey of loss-of-function variants in human protein-coding genes. Science 335, 823–828 (2012).

44. D. L. Remington, Alleles versus mutations: Understanding the evolution of genetic architecture requires a molecular perspective on allelic origins. Evolution 69, 3025–3038 (2015).

45. J. G. Monroe et al., Adaptation to warmer climates by parallel functional evolution of CBF genes in Arabidopsis thaliana. Mol Ecol 25, 3632–3644 (2016).

46. P. J. Flood, A. M. Hancock, The genomic basis of adaptation in plants. Curr Opin Plant Biol 36, 88–94 (2017).

47. A. L. Price et al., Principal components analysis corrects for stratification in genome-wide association studies. Nature genetics 38, 904 (2006).

48. G. A. Fox, Drought and the evolution of flowering time in desert annuals. American Journal of Botany, 1508–1518 (1990).

49. M. Bosse et al., Recent natural selection causes adaptive evolution of an avian polygenic trait. Science 358, 365–368 (2017).

50. D. M. Goodstein et al., Phytozome: a comparative platform for green plant genomics. Nucleic acids research 40, D1178–D1186 (2011).

51. K. Katoh, D. M. Standley, MAFFT multiple sequence alignment software version 7: improvements in performance and usability. Molecular biology and evolution 30, 772–780 (2013).

52. P. L. Ralph, G. Coop, Convergent Evolution During Local Adaptation to Patchy Landscapes. PLoS Genet 11, e1005630 (2015).

53. P. Ralph, G. Coop, Parallel adaptation: one or many waves of advance of an advantageous allele? Genetics 186, 647–668 (2010).

54. R. C. O’Malley, J. R. Ecker, Linking genotype to phenotype using the Arabidopsis unimutant collection. The Plant Journal 61, 928–940 (2010).

55. M. T. Rutter, Y. M. Wieckowski, C. J. Murren, A. E. Strand, Fitness effects of mutation: testing genetic redundancy in Arabidopsis thaliana. J Evol Biol 30, 1124–1135 (2017).

56. J. M. Alonso et al., Genome-wide insertional mutagenesis of Arabidopsis thaliana. Science 301, 653–657 (2003).

